# SARS-CoV-2 RNA quantification using droplet digital RT-PCR

**DOI:** 10.1101/2020.12.21.423898

**Authors:** Natalie N. Kinloch, Gordon Ritchie, Winnie Dong, Kyle D. Cobarrubias, Hanwei Sudderuddin, Tanya Lawson, Nancy Matic, Julio S.G. Montaner, Victor Leung, Marc G. Romney, Christopher F. Lowe, Chanson J. Brumme, Zabrina L. Brumme

## Abstract

Quantitative viral load assays have transformed our understanding of – and ability to manage − viral diseases. They hold similar potential to advance COVID-19 control and prevention, but SARS-CoV-2 viral load tests are not yet widely available. SARS-CoV-2 molecular diagnostic tests, which typically employ real-time reverse transcriptase-polymerase chain reaction (RT-PCR), yield semi-quantitative results only. Reverse transcriptase droplet digital PCR (RT-ddPCR), a technology that partitions each reaction into 20,000 nanolitre-sized droplets prior to amplification, offers an attractive platform for SARS-CoV-2 RNA quantification. We evaluated eight primer/probe sets originally developed for real-time RT-PCR-based SARS-CoV-2 diagnostic tests for use in RT-ddPCR, and identified three (Charité-Berlin E-Sarbeco and Pasteur Institute IP2 and IP4) as the most efficient, precise and sensitive for RT-ddPCR-based SARS-CoV-2 RNA quantification. Analytical efficiency of the E-Sarbeco primer/probe set, for example, was ~83%, while assay precision, as measured by the coefficient of variation, was ~2% at 1000 input copies/reaction. Lower limits of quantification and detection for this primer/probe set were 18.6 and 4.4 input SARS-CoV-2 RNA copies/reaction, respectively. SARS-CoV-2 RNA viral loads in a convenience panel of 48 COVID-19-positive diagnostic specimens spanned a 6.2log_10_ range, confirming substantial viral load variation *in vivo*. We further calibrated RT-ddPCR-derived SARS-CoV-2 E gene copy numbers against cycle threshold (C_t_) values from a commercial real-time RT-PCR diagnostic platform. The resulting log-linear relationship can be used to mathematically derive SARS-CoV-2 RNA copy numbers from C_t_ values, allowing the wealth of available diagnostic test data to be harnessed to address foundational questions in SARS-CoV-2 biology.

## Introduction

Quantitative viral load assays have revolutionized our ability to manage viral diseases (1–6). While not yet widely available for SARS-CoV-2, quantitative assays could advance our understanding of COVID-19 biology and inform infection prevention and control measures (7, 8). Most SARS-CoV-2 molecular diagnostic assays however, which use real-time reverse transcriptase PCR (RT-PCR) to detect one or more SARS-CoV-2 genomic targets using sequence-specific primers coupled with a fluorescent probe, are only semi-quantitative. These tests produce cycle threshold (C_t_) values as readouts, which represent the PCR cycle where the sample began to produce fluorescent signal above background. While each C_t_ value decrement corresponds to a roughly two-fold higher viral load (due to the exponential nature of PCR amplification), C_t_ values cannot be directly interpreted as SARS-CoV-2 viral loads without calibration to a quantitative standard (9). Rather, C_t_ values are interpreted as positive, indeterminate or negative based on assay-specific cutoffs and evolving clinical guidelines. Due to differences in nucleic acid extraction method, viral target and other parameters, C_t_ values are also not directly comparable across assays or technology platforms.

Reverse transcriptase droplet digital PCR (RT-ddPCR) offers an attractive platform for SARS-CoV-2 RNA quantification (10, 11). Like real-time RT-PCR, ddPCR employs target-specific primers coupled with a fluorescent probe, making it relatively straightforward to adapt assays. In ddPCR however, each reaction is fractionated into 20,000 nanolitre-sized droplets prior to massively parallel PCR amplification. At end-point, each droplet is categorized as positive (target present) or negative (target absent), allowing for absolute target quantification using Poisson statistics. This sensitive and versatile technology has been used for mutation detection and copy number determination in the human genome (12), target verification following genome editing (13), and copy number quantification for viral pathogens (14–19). Several real-time RT-PCR SARS-CoV-2-specific primer/probe sets have been used in RT-ddPCR (10, 11, 20–22) with results achieving high sensitivity in some reports (11, 21, 23–25), but few studies have rigorously evaluated SARS-CoV-2-specific primer/probe set performance in RT-ddPCR using RNA as a template. Furthermore, no studies to our knowledge have calibrated SARS-CoV-2 viral loads to diagnostic test C_t_ values. Here, we evaluate eight SARS-CoV-2-specific primer/probe sets originally developed for real-time RT-PCR (26), for use in RT-ddPCR. We also derive a linear equation relating RT-ddPCR-derived SARS-CoV-2 viral loads and real-time RT-PCR-derived C_t_ values for a commercial diagnostic assay, the LightMix^®^ Modular SARS-CoV (COVID19) E-gene assay, allowing conversion of existing COVID-19 diagnostic results to viral loads.

## Materials and Methods

### Primer and Probe Sets

Eight SARS-CoV-2-specific primer/probe sets developed for real-time RT-PCR COVID-19 diagnostic assays (26) were assessed for use in RT-ddPCR (Table 1). These included the Charité-Berlin E gene (‘E-Sarbeco’) set (27), the Pasteur Institute RdRp IP2 and IP4 sets (‘IP2’ and ‘IP4’, respectively) (28), the Chinese Centre for Disease Control ORF and N gene sets (‘China-ORF’ and ‘China-N’, respectively) (29), the Hong Kong University ORF and N gene sets (‘HKU-ORF’ and ‘HKU-N’, respectively) (30), and the US-CDC-N1 set (31).

### SARS-CoV-2 Synthetic RNA standards

RT-ddPCR assays were evaluated using commercial synthetic SARS-CoV-2 RNA standards comprising six non-overlapping 5,000 base fragments of equal quantities encoding the Wuhan-Hu-1 SARS-CoV-2 genome (Control 2, Genbank ID MN908947.3; Twist Biosciences, supplied at approximately 1 million copies/fragment/μl). To avoid degradation, RNA standards were stored at −80°C and thawed only once, immediately before use, to perform the analytical efficiency, precision, analytical sensitivity and dynamic range analyses described herein. Moreover, to mimic nucleic acid composition of a real biological specimen, all assays employing these standards were supplemented with a consistent, physiologically relevant amount of nucleic acid extracted from pooled remnant SARS-CoV-2-negative nasopharyngeal swabs (Supplementary Figure 1). Briefly, pooled viral transport medium was extracted in 1ml aliquots on the BioMerieux NucliSens^®^ EasyMag^®^, eluted in 60μl and re-pooled. The resulting material contained DNA from on average 2,200 human cells/μl (as quantified using human RPP30 DNA copy numbers by ddPCR as described in (32)) and 4,400 human RNAse P copies/μl extract (as quantified by RT-ddPCR as described in (33)), concentrations that are in line with human DNA and RNA levels recovered on nasopharyngeal swabs (32, 33).

### Reverse transcriptase droplet digital PCR (RT-ddPCR) for SARS-CoV-2 quantification

RT-ddPCR reactions were performed by combining relevant SARS-CoV-2 RNA template with target-specific primers and probe (900nM and 250nM, respectively, Integrated DNA Technologies; Table 1), One-Step RT-ddPCR Advanced Kit for Probes Supermix, Reverse Transcriptase and DTT (300nM) (all from BioRad), XhoI restriction enzyme (New England Biolabs), background nucleic acid (for reactions employing synthetic RNA template only, see above) and nuclease free water. Droplets were generated using an Automated Droplet Generator (BioRad) and cycled under primer/probe set-specific conditions (see below and Figure 1). Analysis was performed on a QX200 Droplet Reader (BioRad) using QuantaSoft software (BioRad, version 1.7.4).

**Figure 1:**
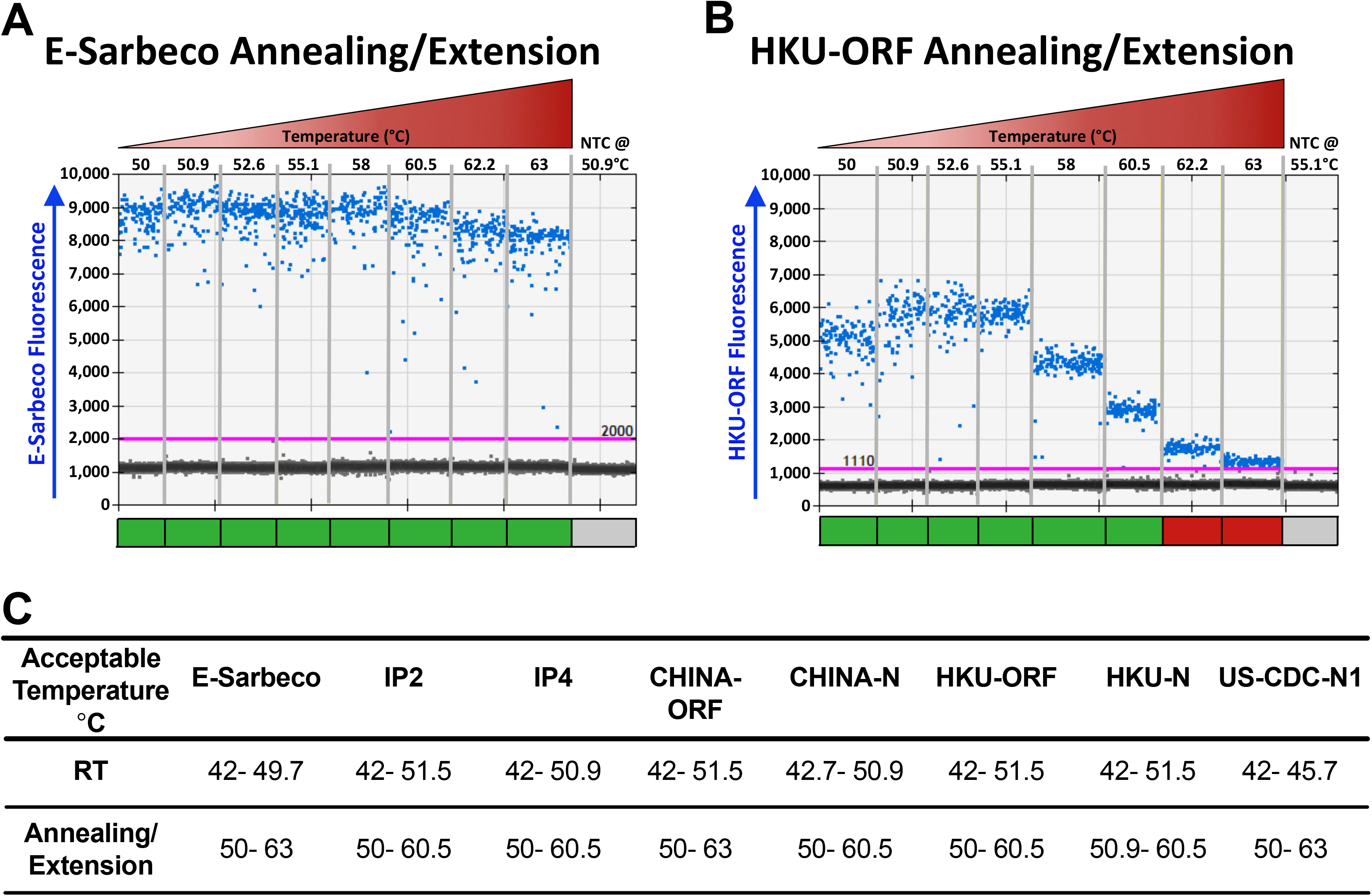
Thermal cycling optimization. (A). RT-ddPCR plots for annealing/extension under a 50-63°C thermal gradient for the E-Sarbeco primer/probe set. A representative RT-ddPCR plot for a no template control (NTC) which only included non-target DNA/RNA (see methods) at the temperature used in subsequent experiments, is also shown. Positive droplets (blue) are above the threshold (pink line); negative droplets (grey) are below the line. Colored boxes below each well indicate if results met standards for inclusion (green) or not (red) (see methods). (B). Same as panel A, but for HKU-ORF primer/probe set. (C). Acceptable RT and annealing/extension temperature ranges for each primer/probe set.

### Thermal cycling temperature optimization

For each primer/probe set, acceptable thermal cycling temperature ranges for reverse transcription (RT) and PCR annealing/extension were determined by modifying the manufacturer-recommended default conditions, which are 42-50°C for 1 hour (for reverse transcription); 95°C for 10 minutes; 40 cycles of (94°C for 30 seconds followed by 50-63°C for 1 minute); 98°C for 10 minutes and 4°C infinite hold. To determine acceptable temperature ranges for reverse transcription, a thermal gradient from 42-51.5°C was performed while fixing the annealing/extension step at 52°C. Using the optimized reverse transcription temperature, a thermal gradient from 50-63°C was then performed to identify acceptable annealing/extension temperature ranges. Temperatures that produced insufficient separation of positive from negative droplets or non-specific amplification were deemed unacceptable, as were those that produced consecutive 95% confidence intervals of copy number estimates outside those of the maximal point-estimate.

### Analytical Efficiency and Precision

The analytical efficiency of each primer/probe set to quantify SARS-CoV-2 RNA by RT-ddPCR was determined using synthetic SARS-CoV-2 RNA standards at 1000 and 100 input copies. A minimum of three (maximum four) technical replicates were performed at each concentration. Analytical efficiency was calculated by dividing the measured SARS-CoV-2 copy number by the expected input copy number, and multiplying by 100. Precision was expressed as the coefficient of variation (CV), expressed as a percentage, across technical replicates.

### Linear Dynamic Range

The linear dynamic range (LDR) of each primer/probe set of interest was determined across a serial 1:2 dilution series from 114,286 to 1.2 SARS-CoV-2 RNA copies/reaction. This range of concentrations was chosen as it crosses the entire range of recommended input copies for a ddPCR reaction seeking to quantify the target of interest (34). Reactions were performed in duplicate. The upper and lower limits of quantification of (ULOQ and LLOQ, respectively) were defined as the upper and lower boundaries of the concentration range over which the relationship between measured and input SARS-CoV-2 RNA copies was linear. This was determined by iteratively restricting the range of concentrations included in the linear regression of measured versus input SARS-CoV-2 RNA copies to identify that which maximized the coefficient of determination (R^2^) value and minimized the residuals.

### Assay Analytical Sensitivity

Assay analytical sensitivity, defined as the Lower Limit of Detection (LLOD), was determined for primer/probe sets of interest by serially diluting synthetic SARS-CoV-2 RNA standards to between 47.6 and 0.74 SARS-CoV-2 RNA copies/reaction. Between 6 and 18 technical replicates were performed for each dilution and results were analyzed using probit regression. The LLOD, determined through interpolation of the probit curve, was defined as the concentration of input SARS-CoV-2 RNA in a reaction where the probability of detection was 95%.

### SARS-CoV-2 RNA quantification in biological specimens, and relationship to C_t_ value

Optimized RT-ddPCR assays were applied to a convenience sample of 48 consecutive remnant SARS-CoV-2-positive diagnostic nasopharyngeal swab specimens that were originally submitted to the St. Paul’s Hospital Virology Laboratory in Vancouver, Canada for diagnostic testing using the Roche cobas® SARS-CoV-2 assay. For these samples, total nucleic acids were re-extracted from 250μl remnant media using the BioMerieux NucliSens® EasyMag® and eluted in 50μl. Eluates were aliquoted and frozen at −80°C prior to single use. SARS-CoV-2 copy numbers were quantified by RT-ddPCR as described above. As our main goal was to characterize the relationship between C_t_ values and SARS-CoV-2 RNA levels without confounding by extraction platform, quantity of input material or SARS-CoV-2 genomic target, we re-tested these extracts using a commercial real-time RT-PCR SARS-CoV-2 diagnostic assay that uses the E-Sarbeco primer/probe set (27): the LightMix® 2019-nCoV real-time RT-PCR assay E-gene target (Tib-Molbiol), implemented on LightCycler 480 (Roche Diagnostics). Finally, to be responsive to a recent recommendation that SARS-CoV-2 viral loads be reported in terms of SARS-CoV-2 RNA copies per human cell equivalents (9), we measured human cells/μl extract by ddPCR as previously described (32) and additionally reported results as SARS-CoV-2 RNA copies/1,000 human cells.

### Statistical Analysis

Statistical analysis was performed using GraphPad Prism (Version 8) or Microsoft Excel (Version 14.7.2).

### Ethical Approval

This study was approved by the Providence Health Care/University of British Columbia and Simon Fraser University Research Ethics Boards under protocol H20-01055.

## Results

### Thermal cycling optimization for SARS-CoV-2 quantification by RT-ddPCR

Eight primer/probe sets originally developed for SARS-CoV-2 diagnostic testing by real time RT-PCR were evaluated for use in RT-ddPCR (Table 1). As these primer/probe sets vary in sequence, amplicon length and SARS-CoV-2 genomic target, we first determined the acceptable temperature ranges for reverse transcription (RT) and PCR annealing/extension. Most primer/probe sets were tolerant to a wide temperature range, and background signal was essentially zero at all temperatures tested (Figure 1). The E-Sarbeco primer/probe set for example produced consistent amplitude profiles, copy number estimates and essentially zero background at annealing/extension temperatures ranging from 50-63°C (Figure 1A and data not shown). The HKU-ORF primer/probe performed acceptably over a 50-60.5°C annealing/extension range, but positive and negative droplet separation was insufficient at higher temperatures (Figure 1B). Acceptable temperature ranges for each primer/probe set are shown in Figure 1C. All subsequent experiments were performed at RT 42.7°C and annealing/extension 50.9°C except those for HKU-ORF and US-CDC-N1, which were performed at RT 45.7°C and annealing/extension 55.1°C as informed by initial qualitative assessments.

### Analytical Efficiency and Precision of SARS-CoV-2 quantification by RT-ddPCR

We next evaluated the analytical efficiency of SARS-CoV-2 RNA quantification for each primer/probe set, calculated as the percentage of input viral RNA copies detected by the assay. We also evaluated precision, calculated as the dispersion of measured copies around the mean (coefficient of variation, CV). Analytical efficiency and precision were evaluated at 1000 and 100 SARS-CoV-2 RNA target input copies. At 1000 input copies, primer/probe set analytical efficiency ranged from 83% (E-Sarbeco) to 15% (US-CDC-N1) (Figure 2A). At 100 copies, the analytical efficiency hierarchy was identical, with values ranging from 74% (E Sarbeco) to 12% (US-CDC-N1). Of all primer/probe sets evaluated, the E-Sarbeco, IP2 and IP4 sets had the highest analytical efficiencies by a substantial margin. At 1000 and 100 target copies, E-Sarbeco analytical efficiency was 83% (95% Total Poisson Confidence Interval [CI]: 79-87%) and 74% (95% CI: 63-84%), respectively; IP2, analytical efficiency was 70% (95% CI: 67-73%) and 55% (95% CI: 46-64%), respectively; and IP4 analytical efficiency was 69% (95% CI: 66-72%) and 59% (95% CI: 50-69%), respectively. In contrast, analytical efficiency of the China-ORF primer/probe set was only 46% and 39% at 1000 and 100 input copies, respectively, and the analytical efficiencies of the remaining sets were less than 30% regardless of input copy number. Furthermore, while measurement precision generally decreased at the lower template concentration (35), the E-Sarbeco, IP2 and IP4 primer/probe sets were nevertheless among the most precise, with coefficients of variation (CV) of less than 5% at 1,000 input copies and less than 15% at 100 input copies (Figure 2B). Combined analytical efficiency and precision data confirmed E-Sarbeco, IP2 and IP4 as the best-performing primer/probe sets in RT-ddPCR (Figures 2C and 2D), so these were moved forward for further characterization.

**Figure 2:**
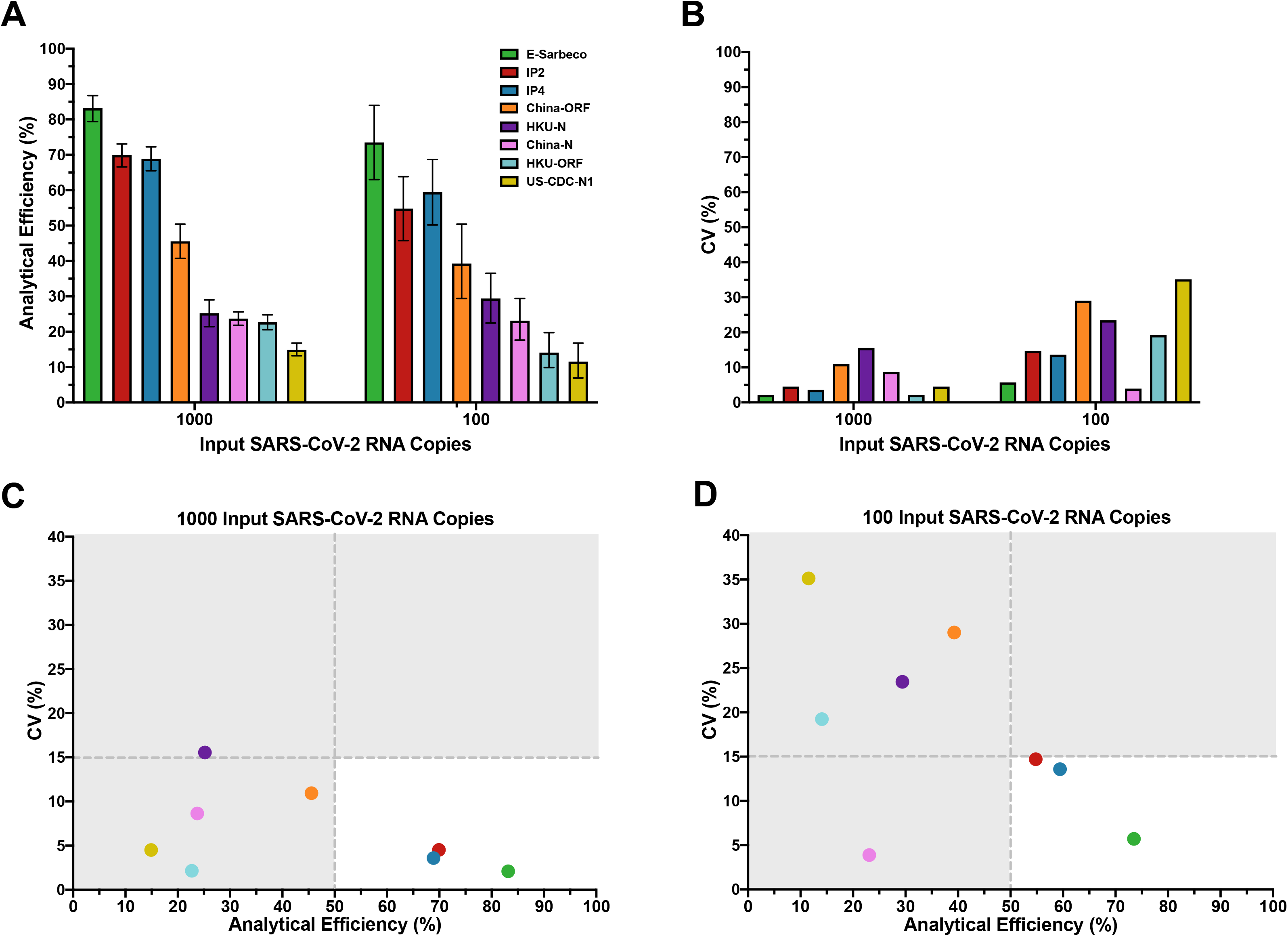
Analytical efficiency and precision of primer/probe sets. (A) Analytical efficiency of each primer/probe set, calculated as the measured divided by the input SARS-CoV-2 RNA copies multiplied by 100%, is shown for reactions containing 1,000 and 100 input copies of synthetic SARS-CoV-2 RNA. Bars represent 95% Total Poisson Confidence Intervals. **(B).** Precision of each primer/probe set, defined as the coefficient of variation (expressed as a percentage, CV%) of measured copies, is shown for reactions containing 1,000 and 100 input copies of synthetic SARS-CoV-2 RNA. (C). Plotting precision versus analytical efficiency at 1,000 input SARS-CoV-2 RNA copies identifies E-Sarbeco, IP2 and IP4 primer/probe sets as having analytical efficiencies >50% and CV (%) <15%. (D). Same as C, but for 100 input SARS-CoV-2 RNA copies.

### Reduced analytical efficiency when IP2 and IP4 are duplexed in RT-ddPCR

As IP2 and IP4 were originally designed for duplexing in real-time RT-PCR (28), we evaluated them in duplex for RT-ddPCR. Duplexing however decreased analytical efficiency, from 70% to 52% (at 1000 input copies) and 55% to 37% (at 100 input copies) for IP2, and from 69% to 49% (at 1000 input copies) and 59% to 38% (at 100 input copies) for IP4 (Supplemental Figure 2A). Duplexing also decreased precision (Supplemental Figure 2B). For IP2, CV increased from 5% to 11% when duplexing at 1000 input copies, and from 15% to 25% when duplexing at 100 input copies. For IP4, CV increased from 4% to 7% (1000 input copies) and from 14% to 21% (100 input copies) with duplexing. Duplexing of these reactions is therefore not recommended in RT-ddPCR, and all IP2 and IP4 assays were performed as single reactions.

### Linear Dynamic Range and Limits of Quantification of SARS-CoV-2 RNA by RT-ddPCR

Droplet digital PCR can achieve absolute target copy number quantification without a standard curve. To investigate the linear dynamic range (LDR) of quantification of the E-Sarbeco, IP2 and IP4 assays, we set up 18 two-fold serial dilutions of synthetic SARS-CoV-2 RNA beginning at 114,286 copies/reaction (this copy number is obtained when 120,000 copies are added to a 21μl reaction, of which 20μl is used for droplet generation) and ending with 2.32 copies/reaction. This input copy number range crosses nearly the entire manufacturer-recommended template input range for ddPCR reactions seeking to quantify the target of interest, which is 1-100,000 copies/reaction (36).

The LDR of each assay was determined by iteratively restricting the range of concentrations included in the linear regression of measured versus input SARS-CoV-2 RNA copies to identify the range that maximized the R^2^ value and minimized the residuals. For E-Sarbeco, the regression spanning 18.6-114,286 input SARS-CoV-2 RNA copies per reaction, an approximately 6,100-fold concentration range, yielded an R^2^ value of 0.9995 (Figure 3A, left). Restricting the linear regression to this range also minimized the residuals of all included data points to ±0.065log_10_ copies/reaction (Figure 3A, right). The IP2 assay, while less efficient than E-Sarbeco, had the same estimated LDR of 18.6-114,286 input copies/reaction (Figure 3B, left). This produced an R^2^ value of 0.9995 and residuals within ±0.065log_10_ copies/reaction across the LDR (Figure 3B, right). The LDR of IP4 was estimated as 37.2-114,286 input copies/reaction, an approximately 3,000-fold range, which yielded an R^2^= 0.9975 and produced residuals within ±0.11log_10_ copies/reaction across this range (Figure 3C). For all three assays, 114,286 input copies/reaction should be considered a conservative estimate of the upper limit of quantification, as saturation of the RT-ddPCR reaction or loss of linearity was still not achieved at this concentration.

**Figure 3:**
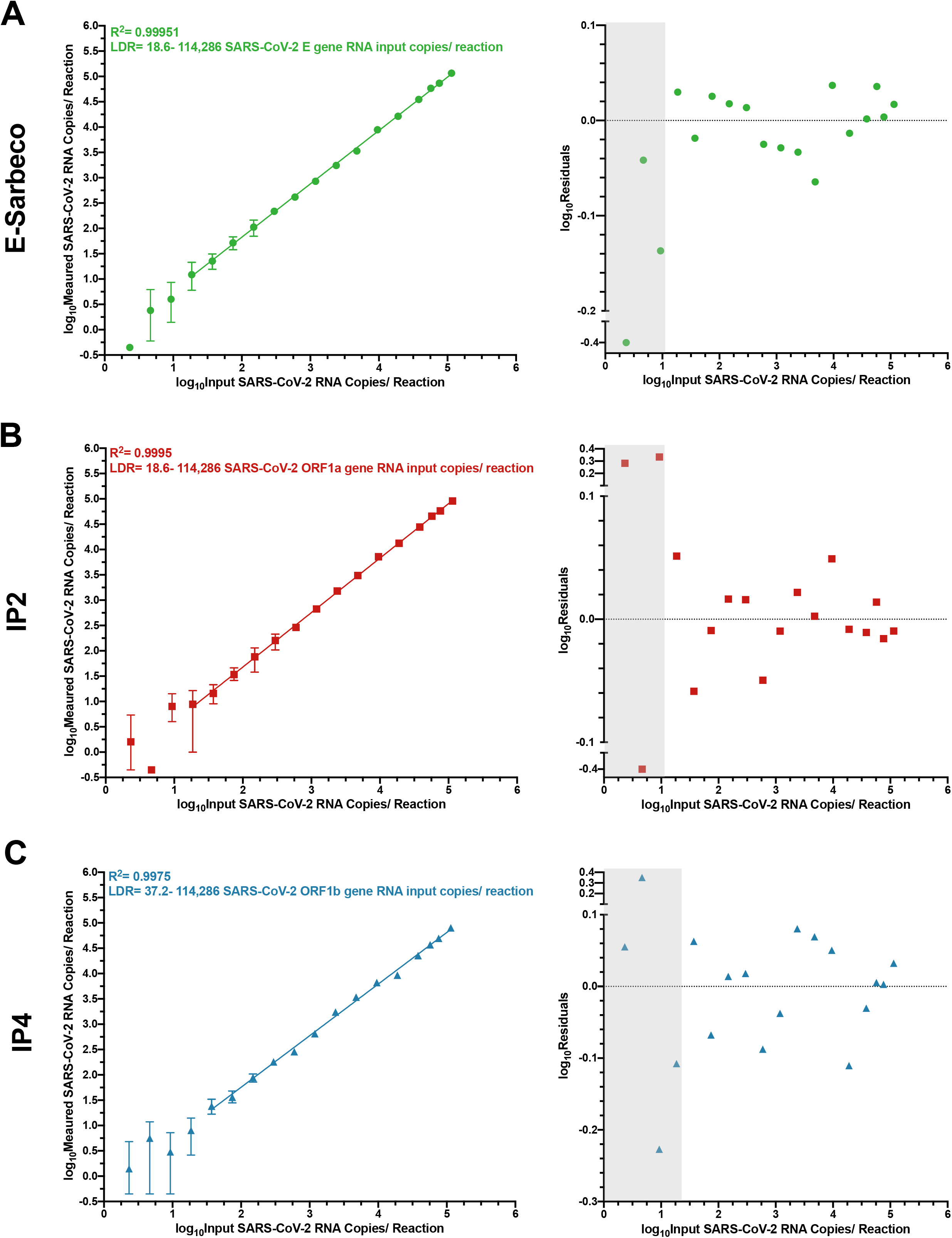
Linear Dynamic Range (LDR) of E-Sarbeco, IP2 and IP4 RT-ddPCR assays. (A). left: log_10_Measured SARS-CoV-2 RNA copies over serial dilutions of synthetic SARS-CoV-2 RNA standards ranging from 114,286 to 2.32 copies/reaction (shown as log_10_ values), using the E-Sarbeco primer/probe set. Error bars indicate 95% Total Poisson Confidence Intervals for two merged replicates, where in some cases error bars are too small to visualize. The regression line joins all data points included in the LDR, where the lower boundary of the LDR represents the lower limit of quantification (LLOQ) of the assay. Data points that yielded undetectable measurements are set arbitrarily to −0.35log_10_Measured copies/reaction for visualization. right: Log_10_Residuals, calculated as log_10_Measured SARS-CoV-2 RNA copies/reaction minus log_10_Calulated SARS-CoV-2 RNA copies/reaction from the LDR regression. Grey shading indicates data points outside the LDR. Residuals for data points that yielded undetectable measurements are arbitrarily set to −0.4 for visualization. (B). Same as A, but for the IP2 primer/probe set (C). Same as A, but for the IP4 primer/probe set.

### Lower Limit of Detection of SARS-CoV-2 RNA by RT-ddPCR

We next determined the lower limit of detection (LLOD) of the E-Sarbeco, IP2 and IP4 RT-ddPCR assays (Figure 4). Probit regression analysis applied to serial dilutions of synthetic SARS-CoV-2 RNA standards revealed the E-Sarbeco RT-ddPCR assay to be the most analytically sensitive of the three, which is consistent with it also having the highest analytical efficiency. Specifically, the estimated LLOD of the E-Sarbeco assay was 4.4 (95% Confidence Interval [CI]: 2.4-5.7) SARS-CoV-2 RNA copies/reaction (Figure 4A). The estimated LLOD of the IP2 assay was 7.8 (95% CI: 4.4-10.3) SARS-CoV-2 RNA copies/reaction (Figure 4B), while that of IP4 was 12.6 (95% CI: 6.9-16.5) SARS-CoV-2 RNA copies per reaction (Figure 4C).

**Figure 4:**
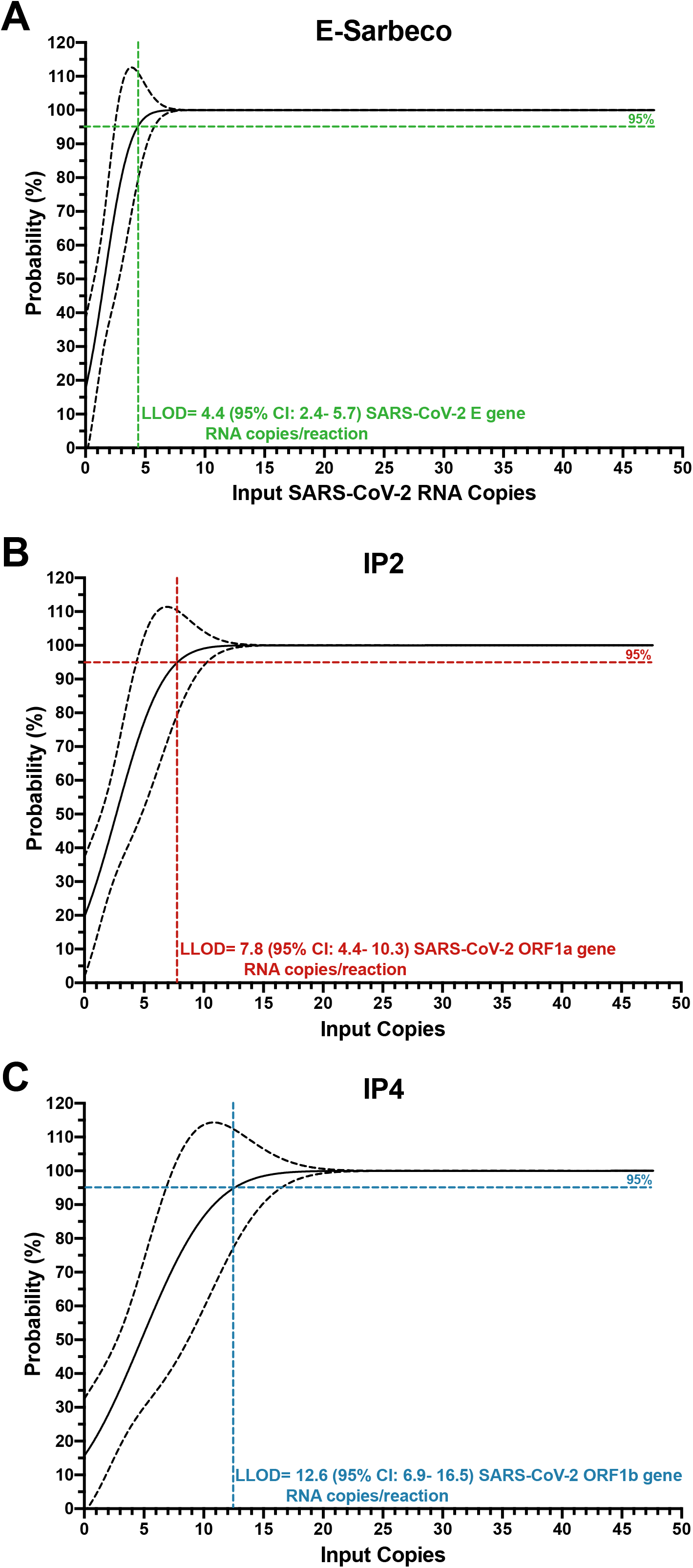
Lower Limit of Detection (LLOD) of the E-Sarbeco, IP2 and IP4 RT-ddPCR assays. (A). The probability of detecting SARS-CoV-2 RNA (%) in 1:2 in serial dilutions of synthetic SARS-CoV-2 RNA from 47.6 to 0.74 input copies/reaction using the E-Sarbeco primer/probe set is analyzed using probit regression (solid black line; dashed line denotes the 95% confidence interval). The LLOD, defined as the concentration of SARS-CoV-2 RNA in a reaction where the probability of detection in the assay was 95%, was interpolated from the standard curve and is shown as a colored dashed line (B). Same as A, but for the IP2 primer/probe set (C). Same as A, but for the IP4 primer/probe set.

### SARS-CoV-2 viral loads in biological samples

SARS-CoV-2 viral loads were measured in 48 confirmed SARS-CoV-2 positive samples using the E-Sarbeco, IP2 and IP4 primer/probe sets (note that samples with original diagnostic test C_t_ values <19 required RNA extracts to be diluted up to 1:200 prior to quantification to ensure that input copies measurements fell within each assay’s LDR). The results revealed that SARS-CoV-2 RNA in these biological samples varied over a 6.2 log_10_ range (Figure 5A). Average copy numbers measured using the E-Sarbeco assay (which targets the E gene) were higher than those using the IP2 and IP4 assays (which target ORF1a and ORF1b, respectively) (Figure 5A). This is consistent with assay analytical efficiency (Figure 2) and *in vivo* coronavirus RNA expression patterns, where transcripts covering the 3’ end of the genome are more abundant than those covering the 5’ end (37–40). Specifically, the median E-gene copy number was 5.1 (IQR 3.9-5.7) log_10_copies/μl extract compared to a median of 4.9 (IQR 3.9-5.5) log_10_copies/μl extract for the IP2 target, and a median of 4.9 (IQR 3.9-5.6) log_10_ copies/μl extract for the IP4 target. SARS-CoV-2 E-gene, IP2 and IP4 copy numbers in biological samples correlated strongly with one another (Spearman’s ρ>0.99; p<0.0001 for all pairwise analyses; Figure 5BCD). Consistent with comparable ORF1a and ORF1b RNA transcript levels *in vivo* (37, 38, 40), IP2 and IP4 copy numbers were also highly concordant (Lin’s concordance correlation coefficient, ρc=0.9996 [95% CI: 0.9993-0.9998]) (Figure 5D). Based on a recent recommendation (9), we also report our results in terms of SARS-CoV-2 RNA copies per human cell equivalents: results for E-Sarbeco spanned an 7-fold range from 1.05 to 7.3 log_10_SARS-CoV-2 RNA copies/1,000 human cells, with IP2 and IP4 log_10_ copy numbers lower, as expected (Supplemental Figure 3A). The Spearman’s correlation between absolute and human cell-normalized viral loads was strong (ρ**=**0.9717; p<0.0001; Supplementary Figure 3B), which is consistent with the assumption that the amount of biological material collected by nasopharyngeal swabs is relatively consistent.

**Figure 5:**
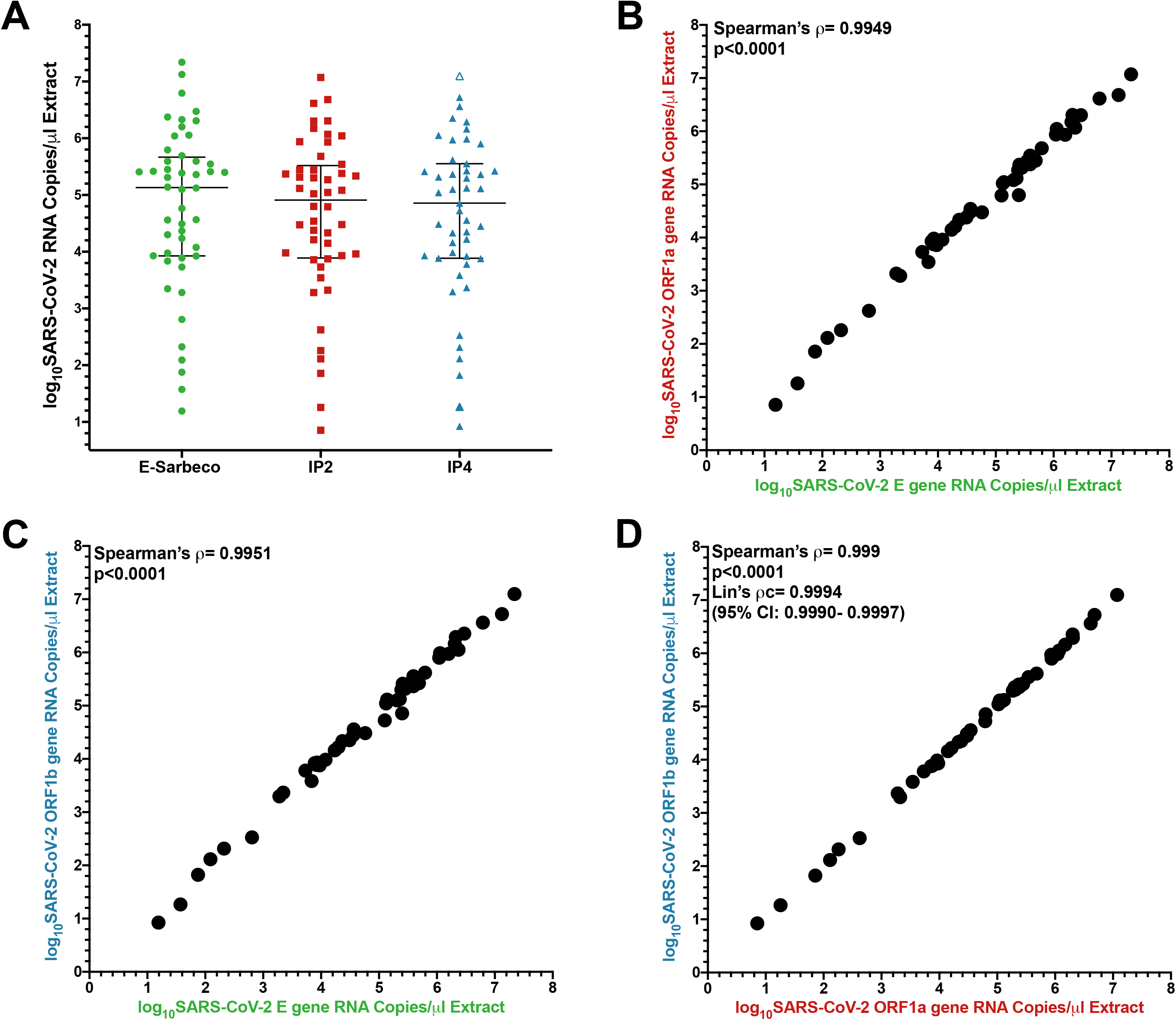
Log_10_SARS-CoV-2 RNA loads in diagnostic specimens. (A). SARS-CoV-2 E (green circles), ORF1a (red squares) and ORF1b (blue triangles) gene copy numbers, expressed as RNA copies/μl of nucleic acid extract. Line and bars indicate median and interquartile range, respectively. **(B)** Correlation between Log_10_SARS-CoV-2 E and ORF1a gene RNA copies/μl extract. (C). Correlation between Log_10_SARS-CoV-2 E and ORF1b gene RNA copies/μl extract. (D) Correlation and Concordance between Log_10_SARS-CoV-2 ORF1a and ORF1b gene RNA copies/μl extract.

### Inferring SARS-CoV-2 viral loads from diagnostic C_t_ values

Finally, we characterized the relationship between C_t_ values produced by a commercial COVID-19 diagnostic platform and SARS-CoV-2 RNA copy numbers. We selected the LightMix® 2019-nCoV real-time RT-PCR assay, E-gene target (Tib-Molbiol), implemented on a LightCycler 480 (Roche Diagnostics) because commercial diagnostic reagents comprising the E-Sarbeco primer/probe set exist for this platform (27) and because it takes purified nucleic acids as input, thereby allowing direct comparison of results from the same starting material (real-time RT-PCR platforms that take biological material as input are suboptimal for such a comparison because the onboard extraction introduces an additional variable). As the C_t_ values reported for the LightMix® assay are based on a 9μl extract input volume, our primary analysis reported RT-ddPCR results in terms of SARS-CoV-2 copies equivalent (*i.e.* SARS-CoV-2 copies in 9μl of extract), to allow direct conversion of C_t_ values to absolute viral copy numbers.

Sample C_t_ values ranged from 11.34-31.18 (median 18.69 [IQR 16.73-22.69]) using the LightMix® assay. The relationship between C_t_ value and SARS-CoV-2 RNA copy numbers was log-linear, with an R^2^ = 0.9990 (Figure 6). Despite this strong relationship, inspection of the residuals nevertheless suggested modest departures from log-linearity at the extremes of the linear range (Supplementary Figure 4). The relationship between C_t_ value and absolute SARS-CoV-2 E-gene copies can thus be given by log_10_SARS-CoV-2 E gene copies equivalent = −0.3038C_t_ +11.7 (Figure 6). That is, a C_t_ value of 20 corresponds to 453,942 (*i.e.* 5.66 log_10_) SARS-CoV-2 RNA copies, while a C_t_ value of 30 corresponds to 416 (*i.e.* 2.62 log_10_) viral copies. This equation also predicts that the C_t_ values corresponding to the LLOQ and LLOD of the E-Sarbeco RT-ddPCR assays are 34.8 and 36.84, respectively. When measured SARS-CoV-2 RNA copy numbers are expressed as human cell-normalized viral loads, the relationship with Ct value is given by log_10_SARS-CoV-2 E gene copies/1,000 human cells = −0.3041C_t_ + 10.8 (Supplemental Figure 5). An extract that yielded a C_t_ value of 20 therefore is estimated to have contained 48,978 (*i.e.* 4.69 log_10_) SARS-CoV-2 RNA copies/1,000 human cells, while one with C_t_ value of 30 is estimated to have contained 45 (i.e. 1.66 log_10_) copies/1,000 human cells

**Figure 6:**
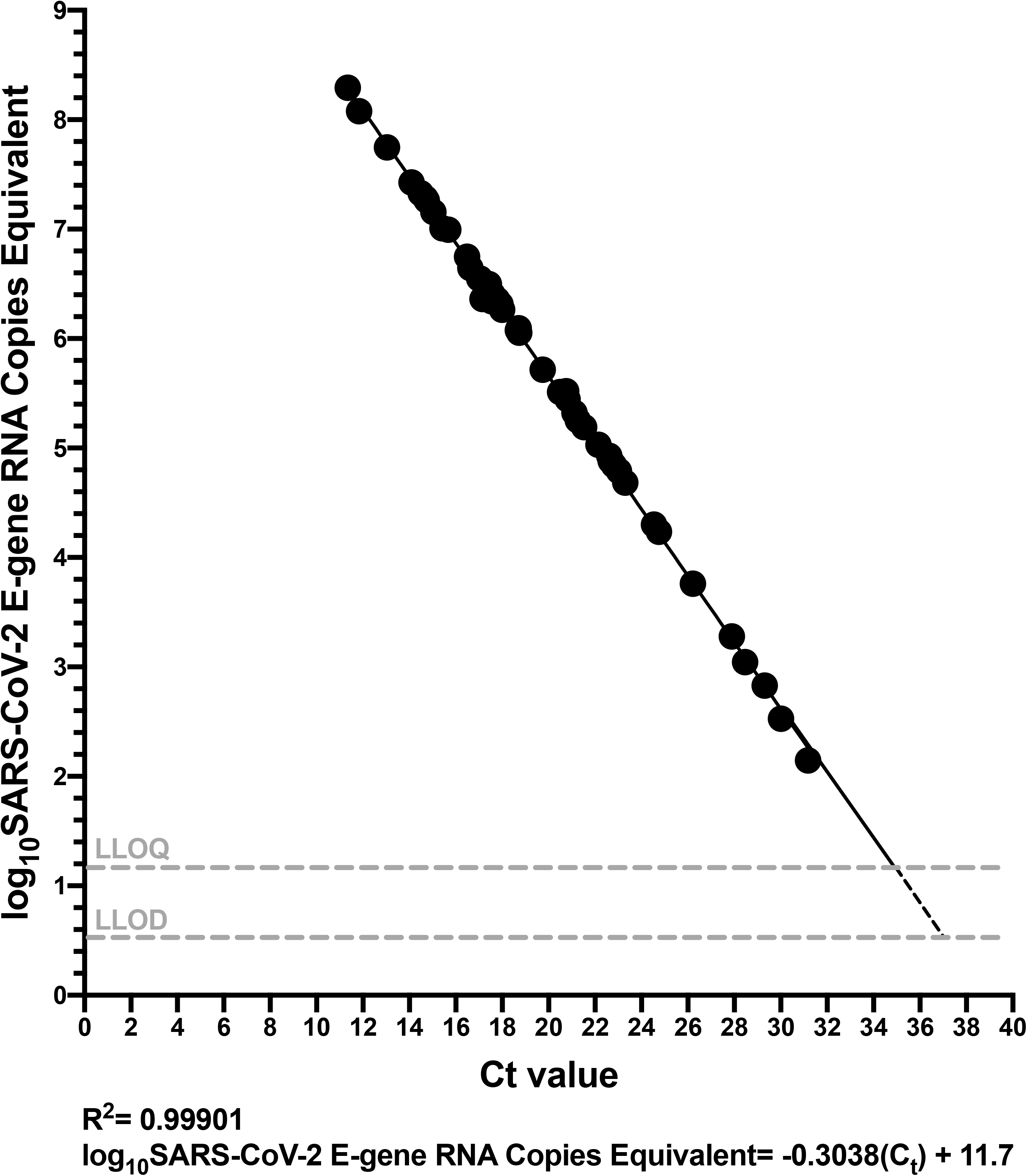
Relationship between SARS-CoV-2 RNA copies equivalent and diagnostic test C_t_ value. C_t_ value, determined using the LightMix® 2019-nCoV real-time RT-PCR assay (E-gene target) is plotted against log_10_SARS-CoV-2 E gene RNA copies equivalent, which represents the number of SARS-CoV-2 RNA copies measured by RT-ddPCR in 9μl extract (the template volume in the LightMix® assay). The linear regression (solid black line) transitions to a dashed line below the LLOQ.

## Discussion

While real-time and droplet digital RT-PCR platforms both employ target-specific primers coupled with fluorescence-based amplicon detection, there are key differences in reaction chemistry (*e.g.* RT-ddPCR reagents must be compatible with water-in-oil droplet partitioning) and probe chemistry (*e.g.* while real-time RT-PCR uses fluorescent quenchers, ddPCR typically uses dark quenchers). As a result, assays developed for one platform may not always translate seamlessly to the other. For example, ddPCR probes should ideally not have a Guanine at their 5’ end because this quenches the fluorescence signal even following hydrolysis (36) but the HKU-N probe has a G at its 5’ end (Table 1).

It is perhaps therefore not surprising that the overall performance of the eight primer/probe sets in RT-ddPCR did not exactly mirror that in real-time RT-PCR (41, 42). Nevertheless, E-Sarbeco, IP2 and IP4, which represented the most efficient and precise primer/probe sets for SARS-CoV-2 RNA quantification by RT-ddPCR are also among the most efficient in real-time RT-PCR (41, 42). Our results also confirm previous reports of the E-Sarbeco primer/probe set performing well in RT-ddPCR (10, 22). Other primer/probe sets however, notably US CDC-N1, HKU-ORF and China-ORF, did not perform as well in our RT-ddPCR assay compared to a previous report (10). One key difference is that, while we used sequence-specific reverse transcription (with the reverse primer) in a one-step RT-ddPCR reaction, the previous study featured an independent reverse transcription reaction primed with random hexamers and oligo dT, which can yield higher efficiency than sequence-specific priming (35, 43–45), to generate cDNA for input into a ddPCR reaction. To our knowledge, ours is the first study to evaluate IP2 or IP4 primer/probe sets in RT-ddPCR.

The analytical sensitivities of the RT-ddPCR assays reported here are nevertheless comparable to existing estimates. The limit of detection of the BioRad SARS-CoV-2 ddPCR Kit (20) is, for example, estimated at 150 copies/mL, which is comparable to our E-Sarbeco RT-ddPCR assay (estimated at 75.8 copies/mL assuming 100% extraction efficiency). Similarly, the LLODs of the TargetingOne (Beijing, China) COVID-19 digital PCR detection kit (23) and a multiplex assay that included the E-Sarbeco primer/probe set (22) were reported at 10 copies/test and 5 copies/reaction, respectively, both comparable to the LLOD determined here. While a number of studies have reported that RT-ddPCR can detect SARS-CoV-2 RNA in low viral load clinical samples with higher sensitivity than real-time RT-PCR (11, 21, 23–25), our study was not designed to evaluate this. Our estimated LLOD of 4.4 copies/reaction by RT-ddPCR using the E-Sarbeco primer/probe set (Figure 4) is in fact comparable to the LLOD reported for many real-time RT-PCR-based COVID-19 diagnostic assays (46).

The ability to quantify SARS-CoV-2 viral loads in biological samples can advance our understanding of COVID-19 biology, and RT-ddPCR offers an attractive platform (7, 8). Our observation that, in a small convenience sample, both absolute and human cell-normalized (9) SARS-CoV-2 loads spanned a more than 6 log_10_ range confirms an enormous viral load range *in vivo* (47) and suggests that some of the high viral load samples measured here were from individuals with early and progressive infection (23, 48–50) or who were experiencing severe disease (7, 8), though clinical information was unknown. Furthermore, our equation relating C_t_ values derived from a commercial diagnostic assay and SARS-CoV-2 RNA copy number means that existing diagnostic test results can be converted to viral loads *without re-testing samples*. While calibration of viral load measurements against all real-time RT-PCR platforms is beyond our scope, this is achievable and in some cases data may already be available (23).

Some limitations merit mention. We only tested eight commonly-used SARS-CoV-2-specific primer/probe sets, and others may exist that adapt well to RT-ddPCR. Our assay performance estimates should be considered approximate, as the manufacturer-reported concentration of the synthetic SARS-CoV-2 RNA standards used in our study may vary by up to 20% error (Twist Bioscience, personal communication). Moreover, we solely evaluated a one-step RT-ddPCR protocol, and therefore assay performance estimates will likely differ from protocols that feature independent cDNA generation followed by ddPCR. We could not precisely define the upper boundary of the linear dynamic range of the E-Sarbeco, IP2 and IP4 RT-ddPCR assays as linearity was maintained at the maximum input of 114,286 target copies/reaction, which already exceeds the manufacturer’s estimated upper range of quantification in a ddPCR reaction (36). Our convenience panel of 48 SARS-CoV-2-positive diagnostic specimens also likely did not capture the full range of biological variation in viral loads, though data from larger cohorts (47) suggests that it was reasonably comprehensive. We also acknowledge that there is measurement uncertainty with real-time RT-PCR C_t_ values that may subtly affect the linear relationship between C_t_ value and RT-ddPCR-derived SARS-CoV-2 viral load described here. Finally, our estimates of assay performance may not completely reflect those of the entire diagnostic process, as the nucleic acid extraction step introduces additional inefficiencies.

In conclusion, primer/probe sets used in real-time RT-PCR-based COVID-19 diagnostic tests can be migrated to RT-ddPCR to achieve SARS-CoV-2 RNA quantification with varying analytical efficiency, precision and sensitivity. Of the primer/probe sets tested, the E-Sarbeco, IP2 and IP4 sets performed best, where LLOQ and LLOD estimates for the E-Sarbeco assay (18.6 and 4.4 copies/reaction, respectively) indicated that RT-ddPCR and real-time RT-PCR have comparable sensitivity. Mathematical inference of SARS-CoV-2 copy numbers from COVID-19 diagnostic test C_t_ values, made possible via the type of calibration performed in the present study, will allow the wealth of existing diagnostic test data to be harnessed to answer foundational questions in SARS-CoV-2 biology.

## Acknowledgements

This work was supported by a COVID-19 rapid response grant from GenomeBC (COVID-115; to ZLB and CFL) and CIHR project grant (PJT-159625; to ZLB). NNK is supported by a Vanier Canada Graduate Scholarship. ZLB holds a Scholar Award from the Michael Smith Foundation for Health Research.

---

**Supplementary Figure 1:**
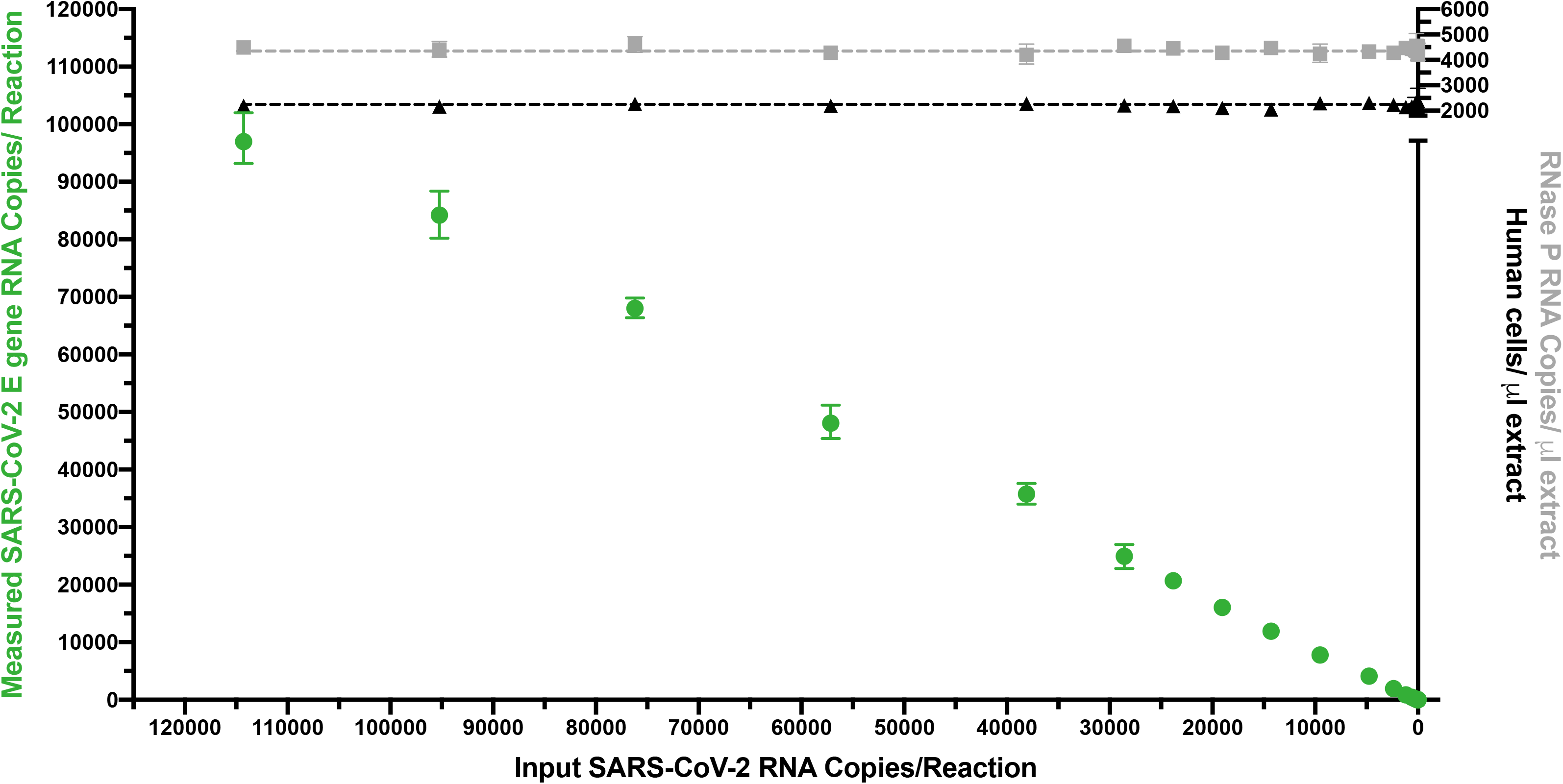
All experiments using synthetic SARS-CoV-2 synthetic standards were performed in a consistent background of human nucleic acids to mimic a real human sample. Example experiment showing consistent levels of background human cells/μl extract (determined by dividing measured human RPP30 DNA copy number by two; black triangles), and human RNAse P RNA levels (grey squares) across a titration of SARS-CoV-2 synthetic RNA standards, measured using the E-Sarbeco primer/probe set (green circles). Error bars indicate 95% Total Poisson Confidence Intervals for two merged replicates, where in some cases error bars are too small to visualize. Grey (RNase P) and black (RPP30) dashed lines indicate copies measured control experiments lacking SARS-CoV-2 RNA.

**Supplementary Figure 2:**
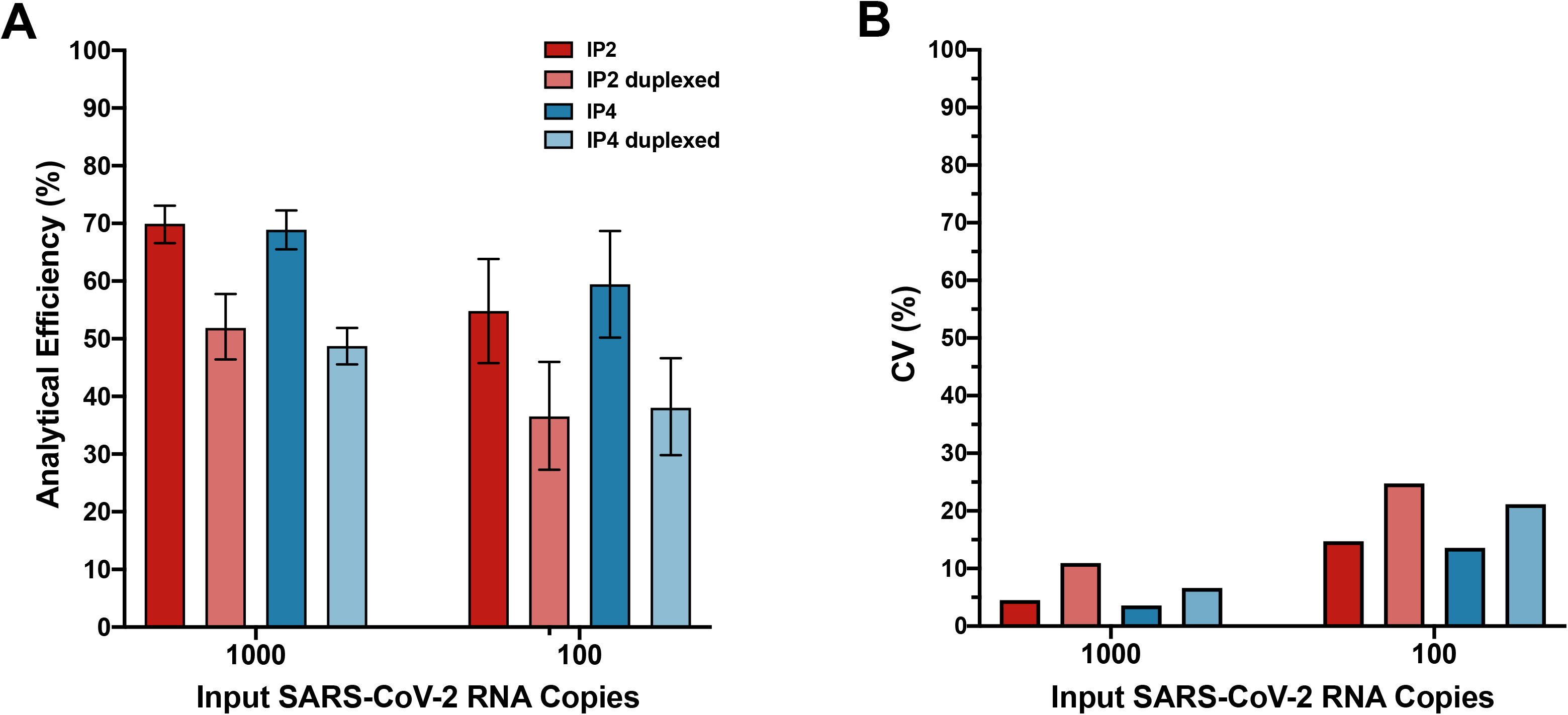
Duplexing the IP2 and IP4 primer/probe sets reduces analytical efficiency and precision. **(A).** Analytical efficiency of SARS-CoV-2 quantification was evaluated for the IP2 and IP4 primer/probe sets when used in separate reactions (dark red and dark blue, respectively) and when duplexed (light red and light blue, respectively), in reactions containing 1,000 and 100 viral RNA input copies. Error bars represent 95% Total Poisson Confidence Intervals. (B). Same as A, but for assay precision (coefficient of variation, CV%).

**Supplementary Figure 3:**
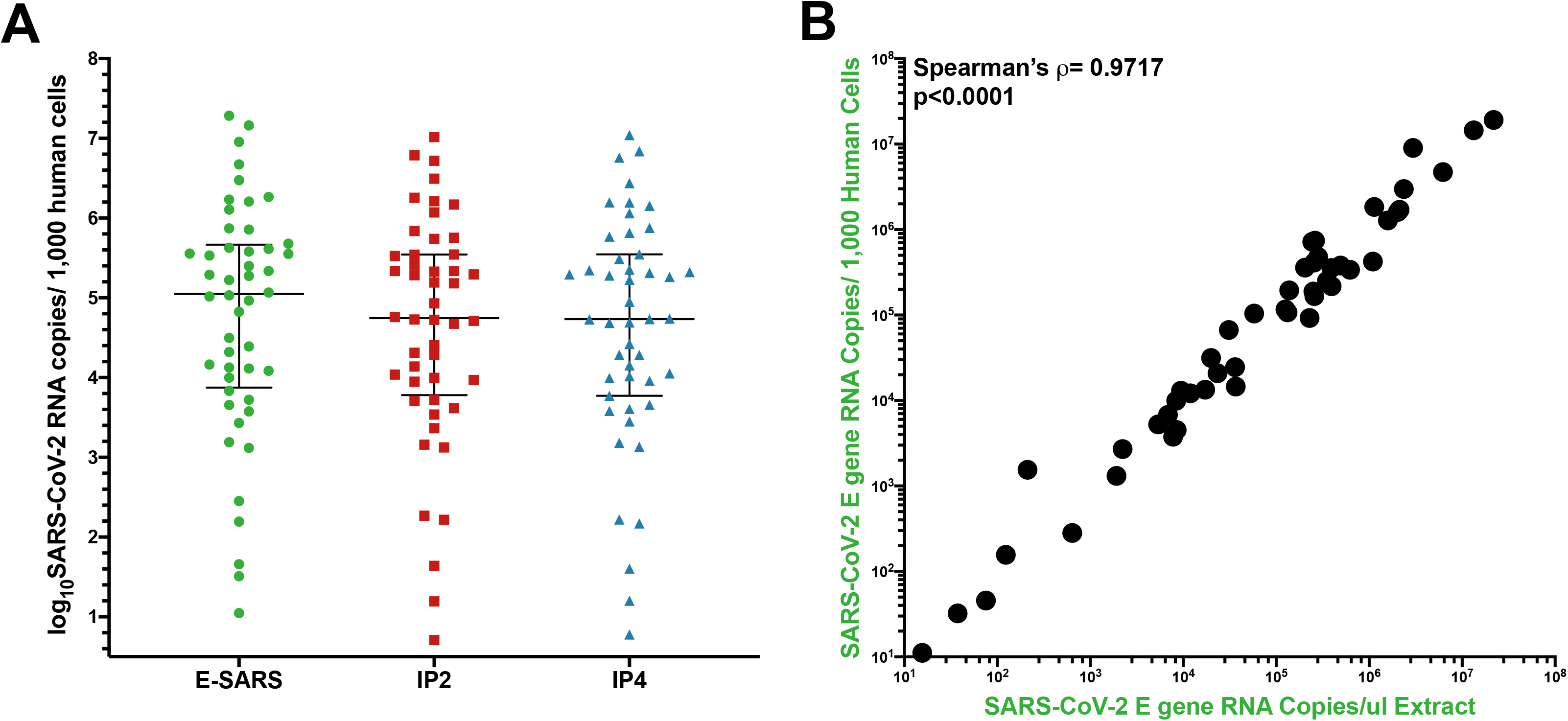
Log_10_SARS-CoV-2 RNA loads in diagnostic specimens, normalized to human cells sampled. **(A)** SARS-CoV-2 E (green circles), ORF1a (red squares) and ORF1b (blue triangles) gene copy numbers, expressed as RNA copies/1,000 human cells. Line and bars indicate median and interquartile range, respectively. (B) Correlation between SARS-CoV-2 RNA copies/μl extract and RNA copies/1,000 human cells.

**Supplementary Figure 4:**
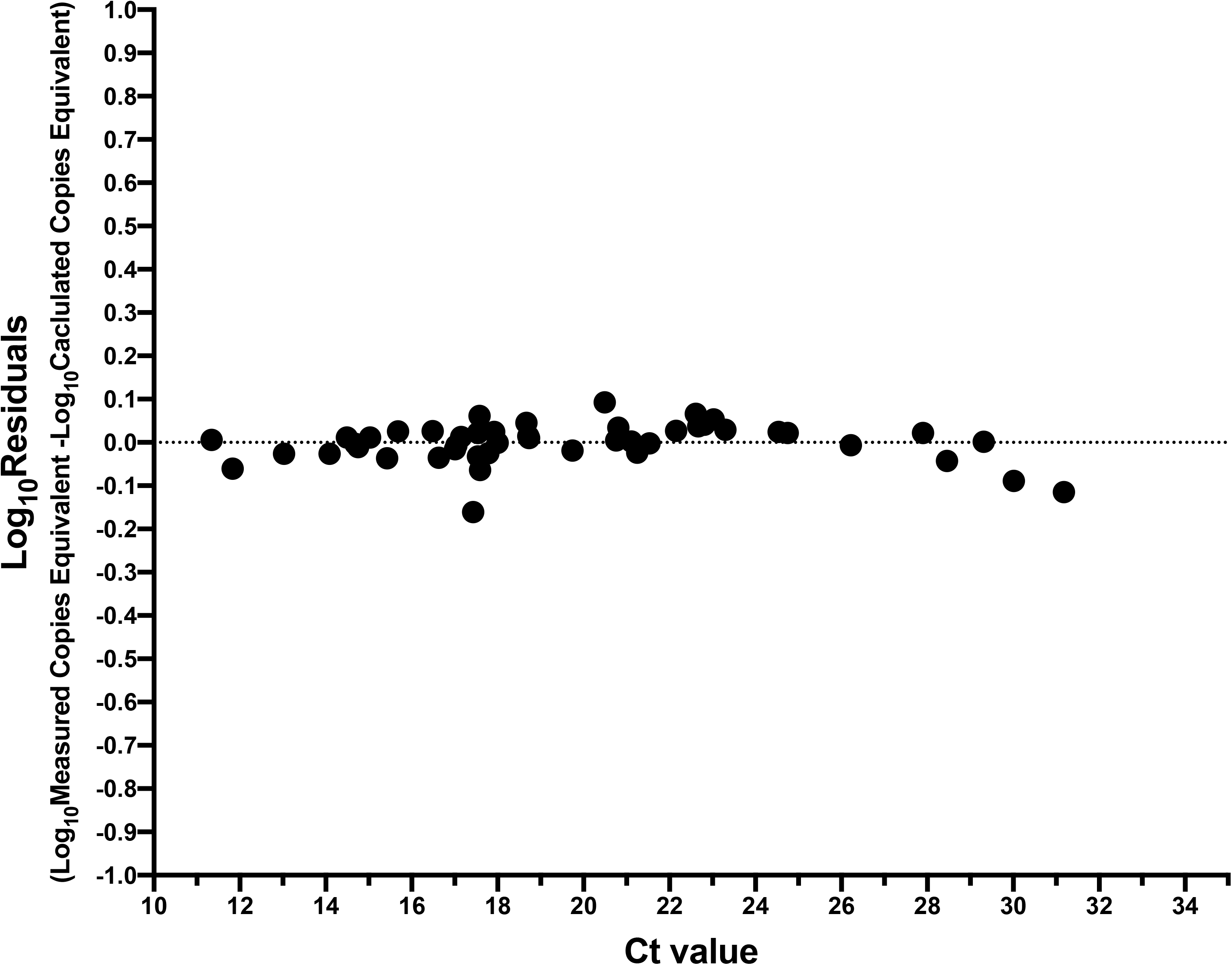
Residuals of relationship between SARS-CoV-2 RNA copies equivalent and diagnostic test C_t_ value. Log_10_Residuals are calculated as log_10_Measured SARS-CoV-2 RNA copies equivalent minus log_10_Calulated SARS-CoV-2 RNA copies equivalent from the regression line shown in Figure 6.

**Supplemental Figure 5:**
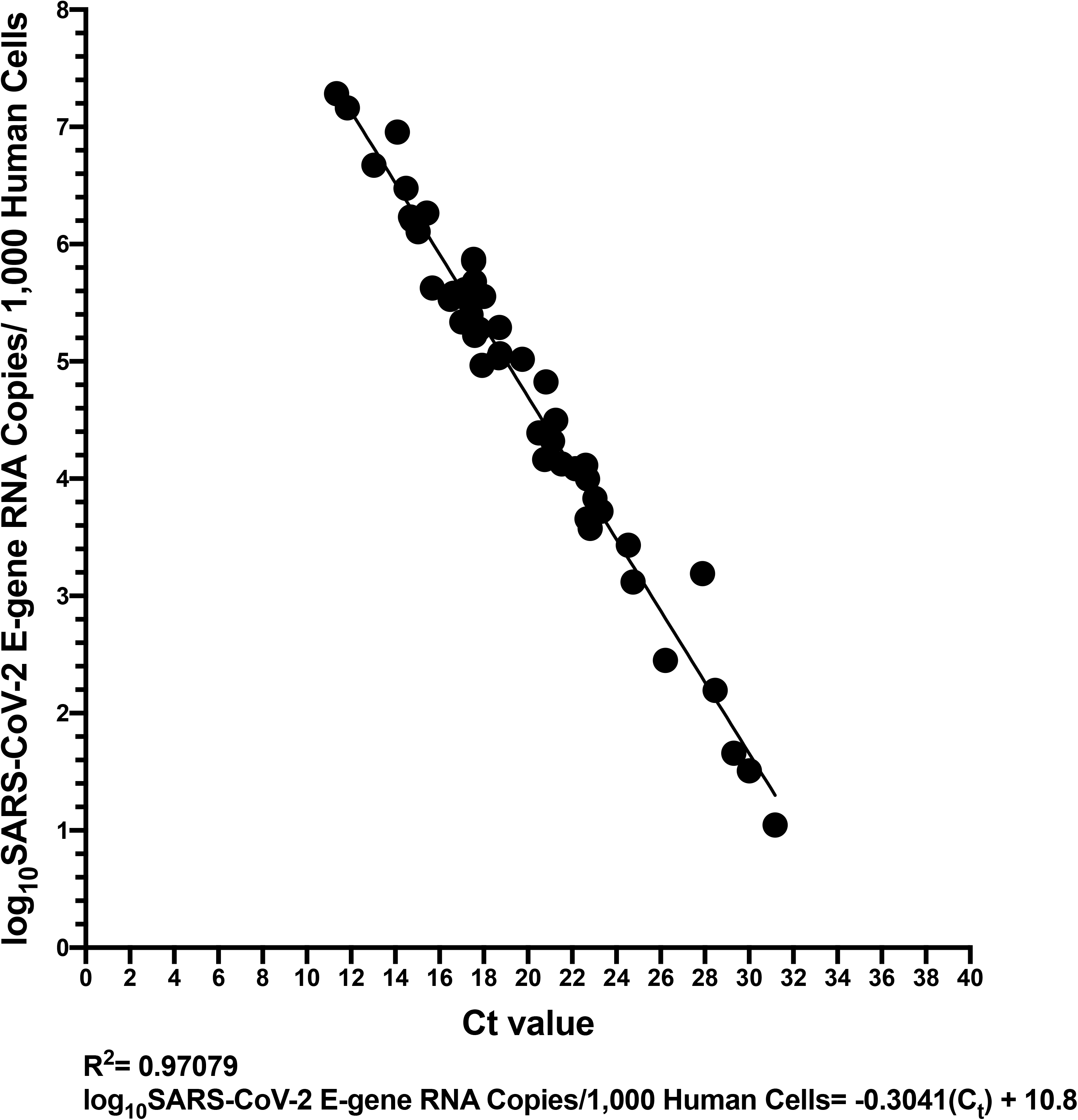
Relationship between SARS-CoV-2 RNA copies/1,000 human cells and C_t_ value. Same data as shown in Figure 6, but where the measured SARS-CoV-2 RNA copies/μl extract were normalized to copies/1,000 human cells. The linear regression is shown as a solid black line.

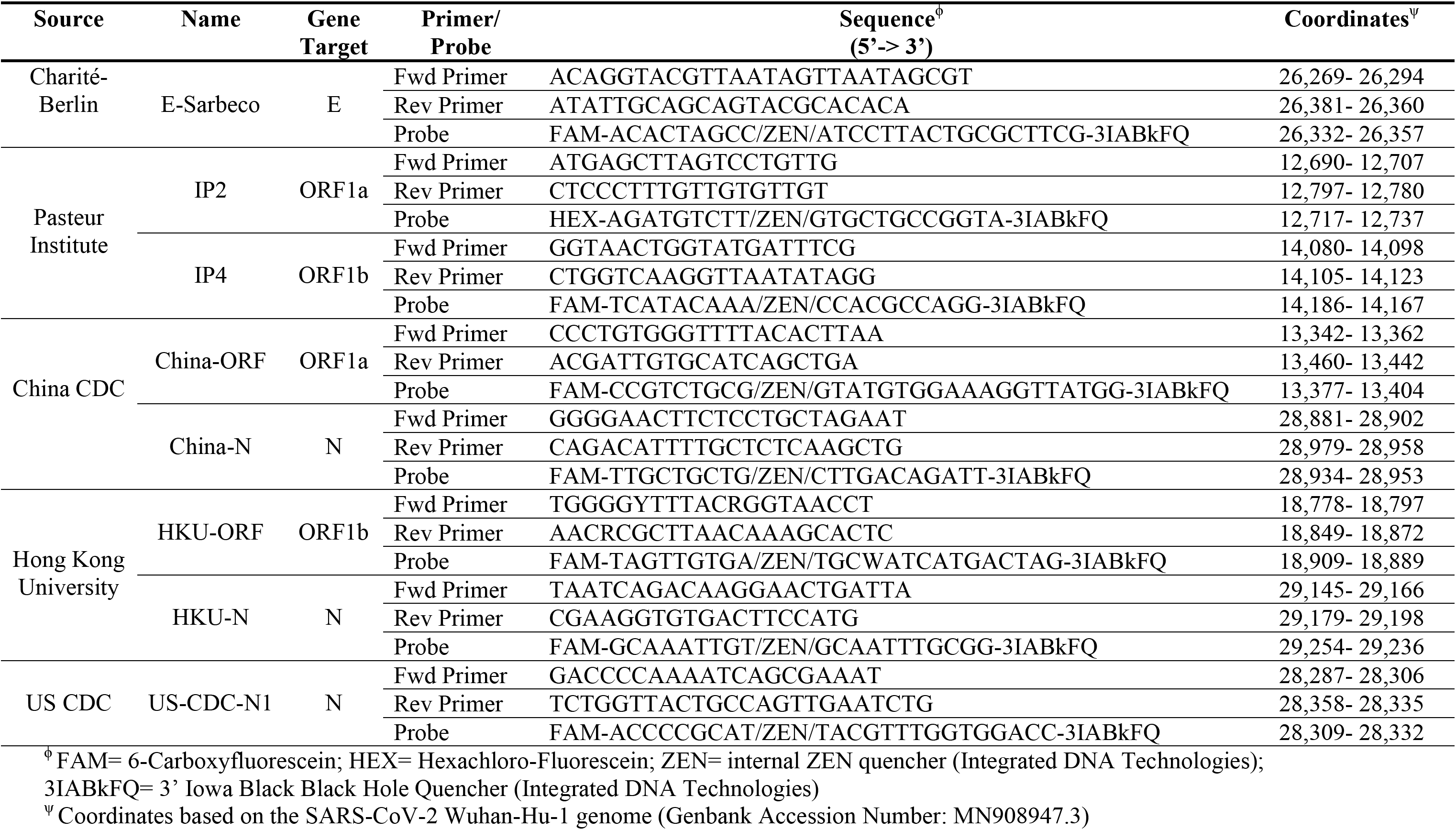

